# Multiple causal DNA variants in a single gene affect gene expression in *trans*

**DOI:** 10.1101/2021.05.05.442834

**Authors:** Sheila Lutz, Krisna Van Dyke, Frank W. Albert

## Abstract

Identifying the specific causal DNA differences in a genome that contribute to variation in phenotypic traits is a key goal of genetic research. *Trans*-acting DNA variants that alter gene expression are important sources of genetic variation. Several genes are known to carry single causal variants that affect the expression of numerous genes in *trans*. Whether these single variants are representative of the architecture of *trans*-acting variation is unknown. Here, we studied the gene *IRA2*, which carries variants with broad, *trans*-acting effects on gene expression in two strains of the yeast *Saccharomyces cerevisiae*. We found that *IRA2* contains at least seven causal nonsynonymous variants. The causal variants were located throughout the gene body and included a pair of neighboring variants with opposing effects that largely canceled each other out. The causal variants showed evidence for non-additive epistatic interactions, in particular among variants at the 5’ end of the gene. These results show that the molecular basis of *trans*-acting variation can involve considerable complexity even within a single gene.

## Introduction

Genetic differences among individuals contribute to variation in phenotypic traits. The identification of causal DNA variants that influence a given trait among the vast majority of variants thought to have no effect is a central challenge in genetics (Rockman, 2012). Quantitative trait locus (QTL) mapping has been used in thousands of studies to identify genomic regions that contain causal variants (Mackay et al., 2009). However, the resulting QTLs are usually broad and span multiple genes and numerous variants. Efforts to identify the molecular cause of a QTL often implicitly assume that the QTL contains only a single causal variant that can be “fine-mapped” through a series of experiments, aided by annotations of gene function and, more recently, computational predictions of variant effect (Adzhubei et al., 2013; Choi and Chan, 2015; Kircher et al., 2014; Sim et al., 2012). Well-known success stories demonstrate that single causal genes and variants can be identified in QTLs (Carneiro et al., 2021; Sutter et al., 2007; Van Laere et al., 2003).

At the same time, the assumption that a typical QTL harbors a single causal variant has frequently turned out to be incorrect (Flint and Mackay, 2009). For example, a single QTL influencing the ability of yeast to grow in high temperature was shown to harbor three different causal genes (Steinmetz et al., 2002). A single chromosome region in *C. elegans* contained multiple independent effects on growth and reproduction (Bernstein et al., 2019). Approximately 40% of QTLs for various physiological traits identified in an outbred rat population were estimated to be due to multiple causal variants (Rat Genome Sequencing and Mapping Consortium, 2013). Some of the most finely dissected QTLs affect traits in the yeast *Saccharomyces cerevisiae* (Fay, 2013). In several yeast QTLs, single genes were found to carry multiple causal variants (Fidalgo et al., 2006; Gerke et al., 2009). Whether typical QTLs tend to be caused by one or multiple variants in one or multiple genes remains an open question, especially given experimental biases that benefit identification of single variants with strong effect compared to multiple variants with smaller effects (Rockman, 2012).

Gene expression variation is a key bridge between DNA variation and organismal phenotypes (Albert and Kruglyak, 2015). In particular, work in yeast crosses (Albert et al., 2018; Brem et al., 2002) and human populations (Grundberg et al., 2012; Wright et al., 2014) revealed important contributions of variants with *trans*-acting effects on gene expression. *Trans*-acting DNA variants change the activity or abundance of a regulatory factor. This factor influences the expression of other genes, which can be located anywhere in the genome. In yeast crosses, *trans*-acting variation arises almost exclusively from “hotspot” regions (Albert et al., 2018) that contain DNA variants that alter the abundance of many genes. *Trans*-acting hotspots are commonly identified by QTL mapping of mRNA abundance as the phenotype of interest. Therefore, like QTLs affecting other traits, the resulting “expression QTLs” (eQTLs) are wide and contain numerous DNA variants.

Due to their broad effects on expression, *trans*-eQTL hotspots have served as models for understanding *trans*-acting regulatory variation. Several yeast hotspots have been dissected to the causal gene (Brion et al., 2013; Lewis and Gasch, 2012; Smith and Kruglyak, 2008) and to single causal variants per gene (Brem et al., 2002; Brown et al., 2008; Fehrmann et al., 2013; Kim et al., 2009; Lutz et al., 2019; Sudarsanam and Cohen, 2014; Yvert et al., 2003; Zhu et al., 2008). Whether these results are representative of most *trans*-eQTL hotspots is unknown.

One particularly prominent *trans*-eQTL hotspot is caused by variation in the *IRA2* gene (Smith and Kruglyak, 2008). *IRA2* encodes a regulator of RAS signaling. Together with *IRA1*, it is one of two paralogous genes that encode the yeast homologs of the human Neurofibromin (NF1) gene, mutations in which can cause neurofibromatosis 1, a disease characterized by uncontrolled cell growth (Ballester et al., 1990; Ratner and Miller, 2015). In a cross between the laboratory strain “BY” and the vineyard isolate “RM”, *IRA2* resides in a *trans*-eQTL hotspot that affects up to 1,240 genes (Albert et al., 2018). Previous work showed that *IRA2* is a causal gene in this hotspot (Smith and Kruglyak, 2008). In addition to mRNA abundance, the *IRA2* locus also affects protein levels for many genes (Albert et al., 2014; Großbach et al., 2019), as well as yeast growth in several different environmental conditions (Bloom et al., 2013; Breunig et al., 2014; Wang and Kruglyak, 2014). Variation in *IRA2* may also underlie QTLs for a variety of traits in strains other than BY and RM (Stojiljkovic et al., 2020; Wang et al., 2019).

In spite of these broad effects on many traits, the causal variant or variants in *IRA2* remain unknown. In part, this is due to the large size of *IRA2*. With a length of 9.24 thousand bases (kb), the protein-coding *IRA2* open reading frame (ORF) is one of the ten longest ORFs in the yeast genome (Cherry et al., 2012), complicating allelic engineering.

Here, we used CRISPR-Swap, an efficient method for yeast genome engineering we recently developed (Lutz et al., 2019), to dissect the molecular basis of the *IRA2* hotspot in the BY / RM yeast cross. We show that the *IRA2* protein coding sequence contains at least seven causal nonsynonymous variants and present evidence that these variants interact with each other in a non-additive, epistatic manner. These results are in contrast to the simpler molecular basis of other *trans*-eQTL hotspots dissected so far, highlighting that even a single causal gene can harbor a complex genetic architecture.

## Results

### Protein-coding variation in *IRA2* affects gene expression in *trans*

The profound *trans* effects of the *IRA2* locus on the mRNA and protein levels of many genes motivated us to identify the nucleotide variant(s) causing these effects. Variants in the coding region of *IRA2* were previously shown to affect the expression of numerous genes (Smith and Kruglyak, 2008). However, the specific coding variants that are responsible for these effects are unknown.

Comparison of the *IRA2* alleles of BY and RM identified 26 nonsynonymous and 61 synonymous variants spread throughout the coding region (**Figure 1A**). Comparison with the *S. cerevisiae* sister species *S. paradoxus* and an *S. cerevisiae* isolate from Taiwan that is presumed to be an evolutionary outgroup to most other strains in this species (Peter et al., 2018) indicates that 19 of the nonsynonymous variants are derived in RM, while 7 are derived in BY (**Table 1**). Causal variants may be expected to occur at sites that are conserved across evolution and encode amino acids with different chemical properties. To ask if any of the 26 nonsynonymous variants are candidates for causing the *IRA2* effect based on these criteria, we analyzed them using the Protein Variation Effect Analyzer (PROVEAN) tool (Choi and Chan, 2015). Four of the derived nonsynonymous variants were marked as “deleterious” (**Table 1**). Variants with strong effects on phenotypes tend to be at low population frequency (Bloom et al., 2019; Fournier et al., 2019). However, only two of the nonsynonymous variants are relatively rare in the population, with a derived allele frequency of 10% or less, and neither of these were predicted to be deleterious (**Table 1**). Together, this information did not pinpoint an individual variant as the obvious source of the effects arising from *IRA2*.

**Figure 1.**
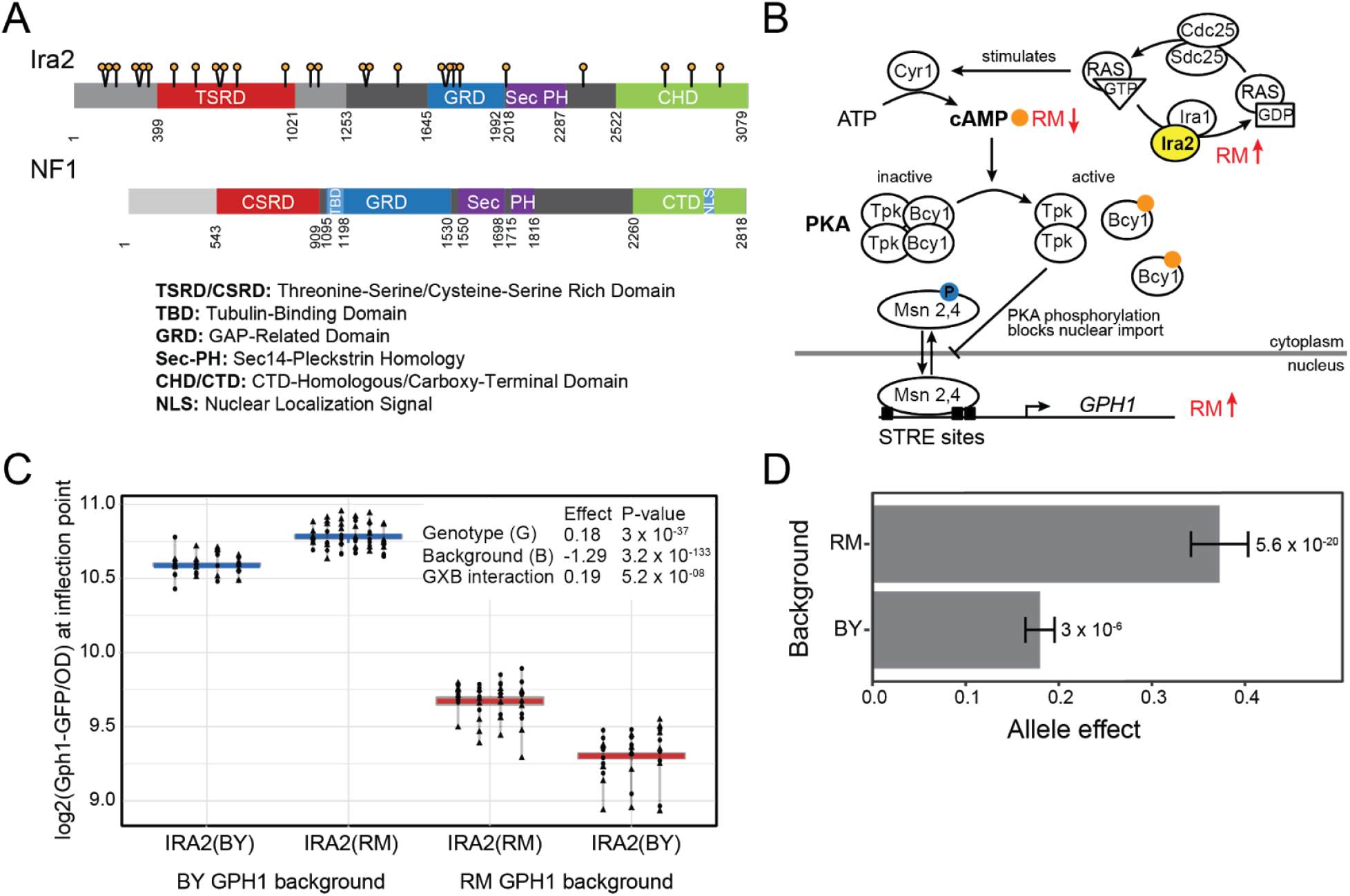
RM variants within the coding region of *IRA2* increase Gph1-GFP expression in *trans*. A: Schematic showing nonsynonymous variants (lollipops) and domains in the Ira2 protein. Defined domains are depicted with colored boxes. The matching dark gray region indicates the continuous homologous region between *IRA2* and NF1. The NF1 schematic is modified from (Bergoug et al., 2020). Details on domain designations for Ira2 can be found in Materials and Methods. B: The role of *IRA2* in the cAMP-PKA pathway including *GPH1*. C: Measurement of the effect of the *IRA2* coding variants in the BY and RM backgrounds. Different shapes represent different plate reader runs. Lines connect measurements of the same transformant. For each strain, 3 – 6 independent transformants were phenotyped 10 – 11 times during two plate reader runs. The boxplots show the median as thicker central lines and the first and third quartiles computed on averaged phenotypes per transformant. D: The *IRA2* allele effect in the BY and RM backgrounds. The allele effect is the log_2_ of the average Gph1-GFP expression in the presence of the RM allele minus that in the presence of the BY allele. Error bars are the standard error of the estimate.

**Table 1.** The 26 nonsynonymous BY/RM variants in *IRA2*

| Variant | Block | Location on chr XV | BY/RM codons<sup>1</sup> | BY/RM amino acids & position | Taiwanese variant | S. *Paradoxus* variant | Derived allele frequency | Derived allele PROVEAN prediction | Derived allele PROVEAN score | RM allele effect | p-value |
| --- | --- | --- | --- | --- | --- | --- | --- | --- | --- | --- | --- |
| 1 | 1 | 171511 | acC/acT | N148H | BY | BY | 0.53 | Neutral | -1.23 | -0.01 | 0.6 |
| 2 | 1 | 171515 | Aac/Cac | S149N | RM | BY | 0.35 | Neutral | -0.105 | -0.017 | 0.35 |
| 3 | 1 | 171671 | aGt/aAt | N201S | BY | BY | 0.18 | Neutral | -1.94 | 0 | 0.99 |
| 4 | 1 | 171973 | Tat/Cat | Y302H | BY | BY | 0.52 | Deleterious | -3.12 | 0.016 | 0.32 |
| 5 | 1 | 171985 | Cat/Gat | H306D | BY | BY | 0.52 | Neutral | 0.3 | -0.01 | 0.58 |
| 6 | 1 | 172102 | Gct/Act | A345T | BY | BY | 0.29 | Neutral | -1.85 | -0.012 | 0.5 |
| 7 | 1 | 172468 | Aat/Gat | N467D | BY | BY | 0.23 | Neutral | -1.24 | -0.011 | 0.57 |
| 8 | 1 | 172768 | Gtc/Atc | V567I | BY | BY | 0.3 | Neutral | -0.07 | 0.008 | 0.6 |
| 9 | 2 | 173080 | Gtg/Atg | V671M | RM | RM | 0.06 | Neutral | -0.15 | 0.009 | 0.66 |
| 10 | 2 | 173105 | aAt/aTt | N679I | BY | BY | 0.5 | Deleterious | -5.64 | -0.027 | 0.17 |
| 11 | 2 | 173340 | caA/caC | Q757H | BY | BY | 0.13 | Neutral | -0.73 | 0.043 | 0.057 |
| 12 | 2 | 174019 | Agc/Tgc | S984C | BY | BY | 0.38 | Deleterious | -2.83 | -0.032 | 0.073 |
| 13 | 2 | 174364 | Ccg/Tcg | P1099S | RM | RM | 0.42 | Neutral | -2.323 | -0.016 | 0.4 |
| 14 | 2 | 174472 | Ctt/Ttt | L1135F | BY | BY | 0.51 | Neutral | -1.85 | 0.049 | **0.0065** |
| 15 | 3 | 175135 | Ata/Gta | I1356V | BY | BY | 0.5 | Neutral | -0.39 | 0.028 | 0.17 |
| 16 | 3 | 175142 | tCt/tTt | S1358F | BY | BY | 0.51 | Neutral | 2.53 | -0.034 | 0.066 |
| 17 | 3 | 175588 | Atc/Gtc | I1507V | RM | RM | 0.36 | Neutral | -0.43 | 0.069 | **0.0027** |
| 18 | 3 | 176239 | Gca/Tca | A1724S | RM | RM | 0.41 | Neutral | -1.057 | -0.034 | 0.06 |
| 19 | 3 | 176268 | aaA/aaT | K1733N | BY | BY | 0.6 | Neutral | 0.56 | 0.026 | 0.163 |
| 20 | 3 | 176348 | aAc/aGc | N1760S | RM | BY | 0.33 | Neutral | 0.653 | -0.007 | 0.69 |
| 21 | 3 | 176441 | tCa/tTa | S1791L | BY | BY | 0.52 | Neutral | -1.65 | -0.003 | 0.89 |
| 22 | 4 | 177067 | Ttt/Gtt | F2000V | BY | BY | 0.39 | Neutral | -0.38 | -0.066 | **0.00047** |
| 23 | 4 | 178159 | Ccc/Tcc | P2364S | BY | BY | 0.41 | Deleterious | -4.08 | 0.042 | **0.02** |
| 24 | 4 | 179291 | gCc/gTc | A2741V | BY | BY | 0.25 | Neutral | 0.88 | 0.006 | 0.74 |
| 25 | 4 | 179652 | ttT/ttG | F2861L | BY | BY | 0.1 | Neutral | 2.56 | 0.006 | 0.74 |
| 26 | 4 | 180056 | aAt/aCt | N2996T | RM | RM | 0.23 | Neutral | 0.072 | 0.009 | 0.62 |

<sup>1</sup>The base in the codon altered by the variant is indicated by a capital letter.

The Ira2 protein negatively regulates the cAMP-PKA signalling pathway by stimulating conversion of RAS GTPase (Ras1,2) from the active GTP-bound form to the inactive GDP-bound form (**Figure 1B**). RAS-GTP activates adenylate cyclase (Cyr1) to produce cAMP. High cAMP results in activation of Protein Kinase A (PKA) and subsequent phosphorylation of its targets (Thevelein and de Winde, 1999). For example, high levels of cAMP reduce the stress response by increasing the phosphorylation and cytoplasmic retention of Msn2,4, which in turn leads to decreased expression of genes containing stress-response elements (STRE) (Görner et al., 1998; Martínez-Pastor et al., 1996). One such gene, *GPH1*, encodes glycogen phosphorylase, an enzyme required for the breakdown of glycogen (Wohler Sunnarborg et al., 2001). The expression of *GPH1* mRNA and Gph1 protein is strongly affected by the *IRA2* locus, with the RM allele resulting in higher expression (eQTL LOD = 21, protein QTL LOD = 60) (Albert et al., 2018, 2014). Therefore, we decided to use *GPH1* tagged with green fluorescent protein (GFP) as a phenotypic readout for experiments aimed at dissecting which variants at the *IRA2* locus affect the expression of other genes in *trans*.

To test if *IRA2* is indeed the causal gene at this locus, we exchanged alleles of the *IRA2* ORF in strains carrying Gph1-GFP using our CRISPR-Swap strategy for rapid genome engineering (Lutz et al., 2019). We then measured Gph1-GFP fluorescence and optical density (OD_600_) of the engineered strains during 24 hours of growth in a plate reader. The Gph1-GFP expression level was extracted at the inflection point of the growth curve and normalized using the OD_600_ value at this time (see Materials and Methods). The presence of the RM allele compared to the BY allele resulted in significantly higher Gph1-GFP expression, confirming that coding variants in *IRA2* contribute to the effects of this locus (**Figure 1C**). The effect was present in both the BY and RM strain backgrounds. However, the effect was larger in RM than in BY (Analysis of variance [ANOVA], p-value for the interaction between *IRA2* genotype and strain background: 5 × 10^−8^; **Figure 1C and D; and Table S3**), suggesting the presence of epistatic interactions between variation in the *IRA2* coding region and other variants elsewhere in the genome.

Increased expression of Gph1-GFP in the presence of the *IRA2*-RM allele is consistent with the direction of known effects of this locus on *GPH1* mRNA and protein (Albert et al., 2018, 2014). Based on the function of Ira2 (**Figure 1B**), increased *GPH1* expression is expected in the presence of a more active *IRA2* allele. Thus, in accordance with previous results (Smith and Kruglyak, 2008), our findings suggest that the RM allele of *IRA2* has higher activity than the BY allele.

### *IRA2* harbors multiple causal variants that show epistatic interactions

To narrow in on the causal variant in the large *IRA2* gene, we divided the *IRA2* ORF into four blocks balancing size and number of nonsynonymous variants (**Figure 2A**). We performed these experiments in BY due to the higher efficiency of genome engineering in this strain background (Lutz et al., 2019). We created 16 chimeric alleles representing all combinations of the four blocks to also allow testing for non-additive (“epistatic”) interactions between the blocks. Each block combination was represented by five or six independent transformants that were each assayed five times for their effect on Gph1-GFP expression.

**Figure 2:**
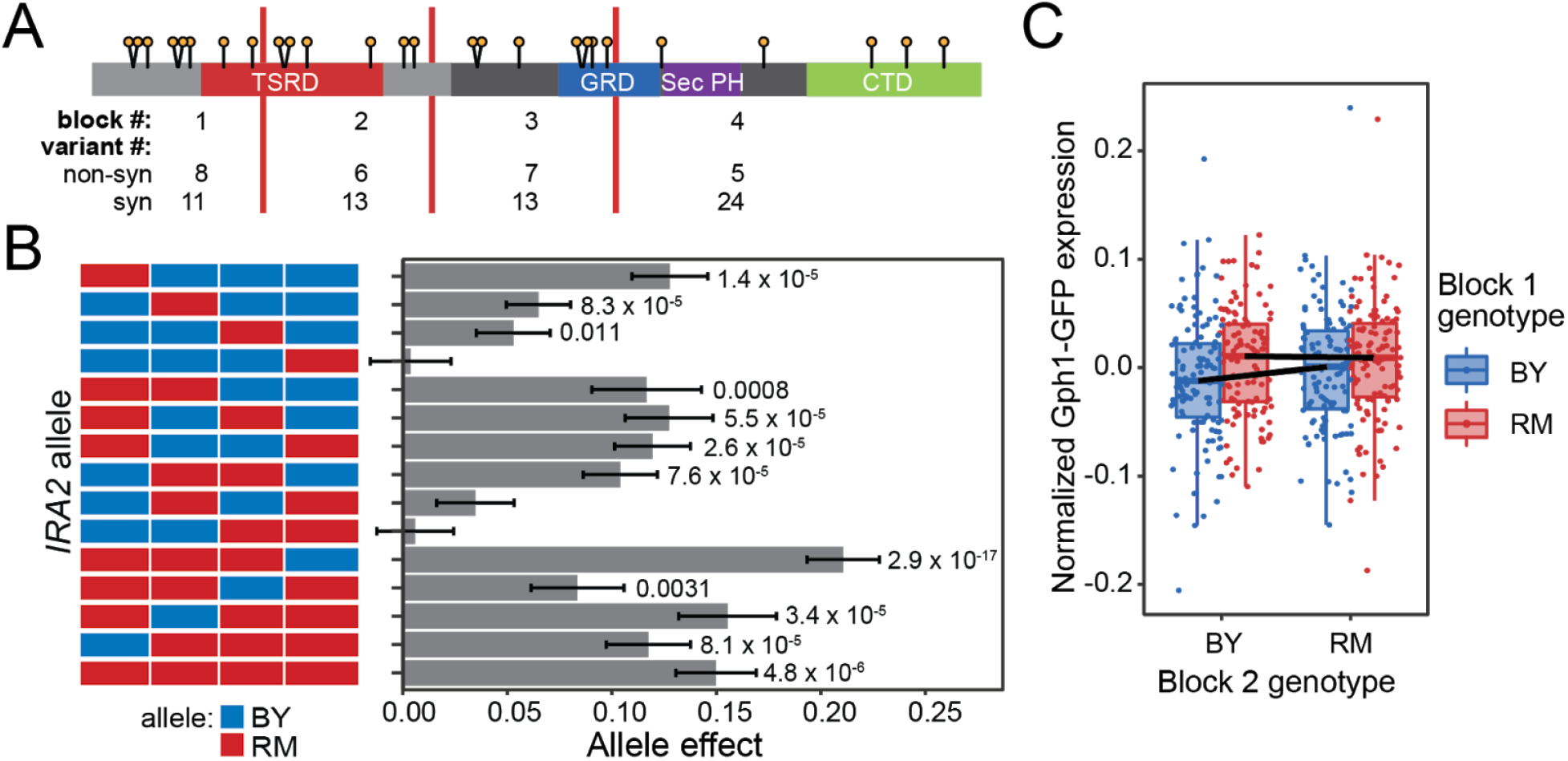
Effect of blocks of *IRA2* variants on Gph1-GFP expression. A: Schematic of Ira2 showing the nonsynonymous variants (lollipops) and block junctions (red vertical lines). B: Allele effect of each of the sixteen block combinations compared to an allele with BY in all blocks. P-values are for a comparison between the given allele and the BY allele, only significant values are displayed. Error bars are standard errors of the given estimate. C: Epistasis between blocks 1 and 2. The figure shows residual Gph1-GFP expression after removal of random effects of plate and transformant identity (Materials and Methods). Measurements are grouped by their alleles at blocks 1 and 2. The remaining two blocks are any combination of BY or RM. The horizontal black lines connect the average allele effect for the indicated genotype. Epistasis is highlighted by the steeper slope of the line when block 1 is BY rather than RM.

At each of the first three blocks, the *IRA2* RM sequence significantly increased Gph1-GFP expression, with the largest effect resulting from block 1 (**Figure 2B; Table 2; Figure S1 and Table S4**). Thus, there must be at least three causal variants in the *IRA2* RM allele, with at least one variant in each of blocks 1, 2 and 3.

**Table 2.** ANOVA of the effects of blocks 1 to 4 and all possible interaction terms

|  | Effect | 95% effect confidence interval | F-value | p-value |
| --- | --- | --- | --- | --- |
| **b1** | 0.13 | 0.08 – 0.19 | 85.57 | **3.9 x 10<sup>-14</sup>** |
| **b2** | 0.06 | 0.01 – 0.12 | 16.86 | **9.9 x 10<sup>-5</sup>** |
| **b3** | 0.05 | 0.001 – 0.11 | 21.93 | **1.2 x 10<sup>-5</sup>** |
| **b4** | 0.004 | -0.05 – 0.06 | 2.96 | 0.09 |
| **b1:b2** | -0.08 | -0.16 – -0.004 | 8.51 | **0.005** |
| **b1:b3** | -0.05 | -0.13 – 0.03 | 0.04 | 0.84 |
| **b1:b4** | -0.01 | -0.10 – 0.07 | 0.04 | 0.85 |
| **b2:b3** | -0.01 | -0.08 – 0.05 | 6.34 | **0.01** |
| **b2:b4** | -0.03 | -0.11 – 0.05 | 1.16 | 0.28 |
| **b3:b4** | -0.005 | -0.12 – 0.03 | 0.00 | 0.97 |
| **b1:b2:b3** | 0.1 | -0.002 – 0.22 | 0.66 | 0.42 |
| **b1:b2:b4** | 0.009 | -0.11 – 0.12 | 3.36 | 0.07 |
| **b1:b3:b4** | 0.09 | -0.03 – 0.20 | 0.05 | 0.82 |
| **b2:b3:b4** | 0.09 | -0.004 – 0.19 | 0.18 | 0.67 |
| **b1:b2:b3:b4** | -0.16 | -0.32 – 0.0002 | 4.22 | **0.04** |

*Effect estimates are polarized such that positive values correspond to higher expression linked* *to the respective RM allele. Effects were computed using a mixed-effects linear model and* *p-values computed by type I ANOVA. Effect size confidence intervals were estimated using a* *bootstrap procedure. The displayed p-values were not corrected for multiple testing. P-values* *less than 0.05 are shown in bold.*

We noticed that one combination of blocks (with the RM at the first 3 blocks and the BY allele at the last block, “RRRB”) resulted in significantly higher Gph1-GFP expression than the allele carrying RM alleles at all four blocks (One-way ANOVA: p = 0.035, **Figure 2B**). Thus, higher Gph1-GFP levels than those driven by the full RM allele can be achieved by combinations of BY and RM variants. Considering that block 4 had no effect when swapped in isolation (**Figure 2B; Table 2 and Table S4**), this difference between the RRRB construct and the full RM allele also suggests that there are epistatic interactions between variants in block 4 and variants in the other blocks.

To comprehensively test for epistatic interactions among blocks, we performed ANOVA on our fully crossed set of block chimeras. Three combinations of blocks showed nominally significant non-additive interactions (**Table 2**), which exceeds the expectation of 0.6 out of 12 interaction tests under a null model without epistasis. The strongest interaction was seen between block 1 and block 2 and is visualized in **Figure 2C**. In all three significant interactions, the observed Gph1-GFP expression driven by the combined blocks was less than that expected from the sum of the individual block effects (note negative effect signs in **Table 2**), indicating negative epistasis between these blocks of variants. In summary, *IRA2* harbors multiple variants that affect Gph1-GFP expression in *trans*, some of which interact in a non-additive manner.

### The effect of each nonsynonymous RM variant is small compared to the full RM allele

To examine the effects of single variants in *IRA2* on Gph1-GFP expression, we engineered 26 *IRA2* strains. Each strain carried the RM allele at a single nonsynonymous variant resulting in a single amino acid change, with all other variants carrying the BY allele. We measured the effect of each single variant allele on Gph1-GFP expression **(Figure 3A; Figure S2; and Table S5)**.

**Figure 3:**
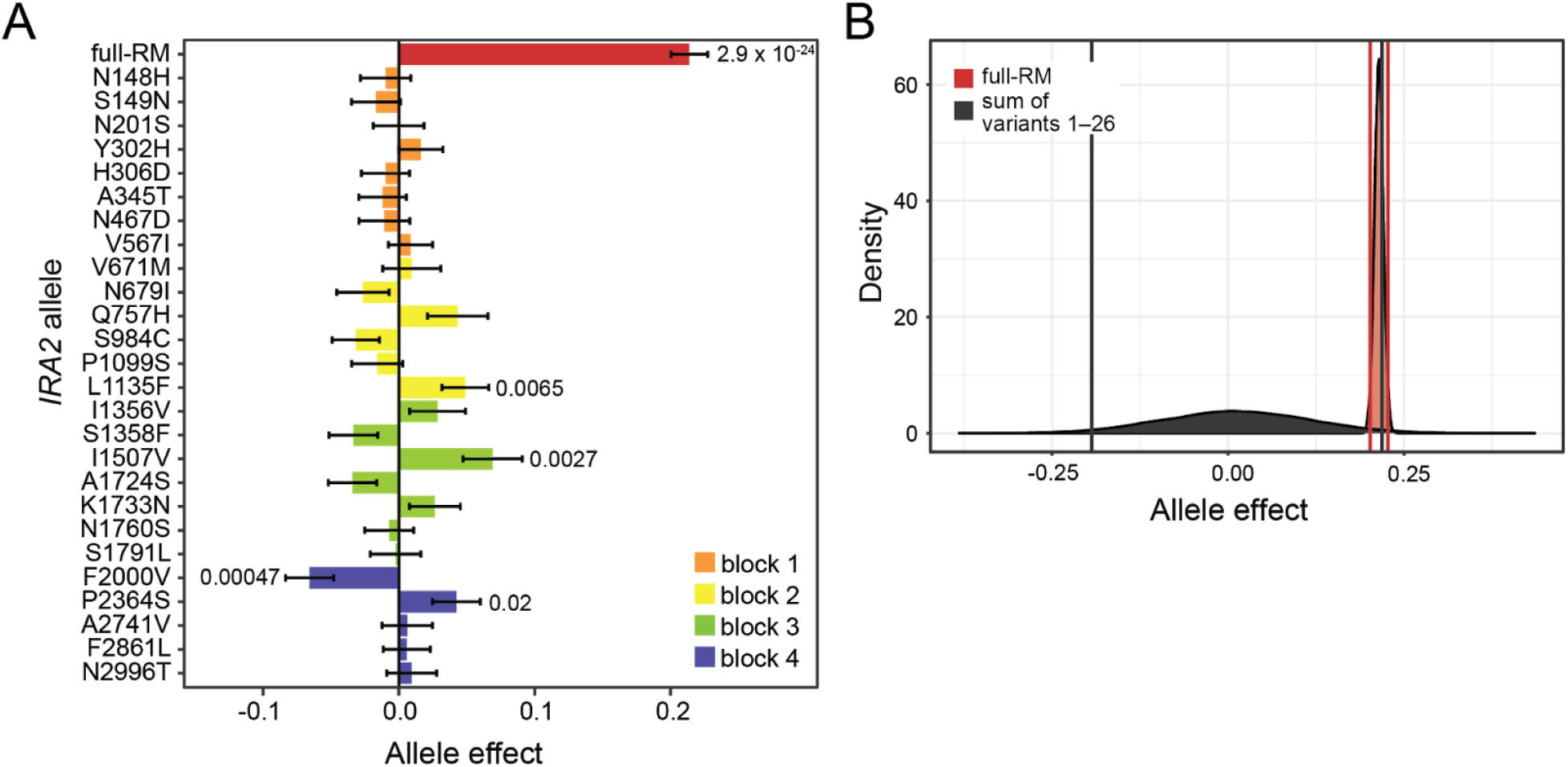
Analysis of the effects of the nonsynonymous variants in *IRA2*. A: Effect of each nonsynonymous RM variant allele on Gph1-GFP expression. Each strain was represented by nine transformants that were each measured five times. The first eight variants (N148H through V567I) were measured three additional times in an additional plate configuration. P-values are shown for a comparison between the given allele and the BY allele. Only significant p-values less than 0.05 are displayed. B: Bootstrap analysis of the effect of the full RM allele compared to the summed effects of the twenty-six variants. The same-colored vertical lines mark the central 95% quantile range for each bootstrap distribution.

Four of the single-variant RM alleles (L1135F, I1507V, F2000V, and P2364S) resulted in Gph1-GFP expression significantly different from the BY allele. Three of these causal variants were predicted to have a neutral effect by PROVEAN, while the fourth, P2364S, was predicted to be deleterious (**Table 1**). None of the four variants were rare in the population, with derived allele frequencies ranging from 0.36 to 0.51. The causal variants are located in three different blocks: L1135F is in block 2, I1507V is in block 3, and F2000V and P2364S are in block 4. Alignment of *IRA2* with its human homolog NF1 revealed that L1135F is in a region that does not have homology with NF1, while I1507, F2000V and P2364S are in regions that are conserved with humans (**Figure 1**). While three of the single RM variants increased Gph1-GFP expression, the F2000V variant reduced it. This direction of effect of F2000V is in the opposite direction of that of the full *IRA2*-RM allele, revealing transgressive segregation of causal variants in a single gene. The effect of F2000V is likely canceled out by its neighboring variant P2364S, at which the RM allele significantly increased Gph1-GFP. The canceling of these variant effects is in agreement with the fact that block 4 as a whole did not significantly alter Gph1-GFP expression (**Figure 2B**). These results show that no single variant underlies the *trans* effect of *IRA2*-RM on Gph1-GFP expression. Instead, a combination of multiple variants must underlie the overall effect of the full RM allele at *IRA2*.

The effects of the single variants were small compared to the effect of the entire *IRA2*-RM allele (**Figure 3A**). To test whether their joint effect can account for the entire *IRA2* effect in an additive manner, we used a bootstrap strategy that accounted for measurement error. We created 10,000 bootstrap datasets by randomly sampling with replacement from our individual measurements. In each dataset, we computed the observed difference between Gph1-GFP abundance for the *IRA2*-BY and the full *IRA2*-RM allele, as well as the sum of the effects driven by each of the 26 single variants. The central 95% of the resulting bootstrap distributions overlapped, such that we cannot formally rule out that the sum of the single variant effects accounts for the effect of the full RM allele in an additive manner (**Figure 3B**). However, the overlap was mostly due to the very long tail of the summed single variant distribution, such that only 3.6% of this distribution exceeded the 2.5% quantile of the full RM allele effect. As also suggested by the significant epistatic interactions among the four blocks above, it remains possible that epistatic interactions among variants across the length of *IRA2* are required to generate the effect of the full *IRA2*-RM allele.

### Multiple, epistatic causal variants are near the 5’-end of *IRA2*

The results above suggest considerable complexity of genetic variation within *IRA2*, with multiple causal variants and non-additive interactions between different regions of the ORF. To dissect this architecture further, we focused on block 1. Although block 1 had the largest effect on Gph1-GFP expression, none of the eight single variants in this block had a significant effect on their own. The variants in block 1 could have real but subtle effects, such that their combined effect could account for the block 1 effect even if no individual variant reached statistical significance. However, bootstrap analysis showed that the additive effects of these variants cannot account for the effect of block 1 (**Figure 4A**), suggesting a non-additive interaction between at least two of the eight variants within block 1.

**Figure 4:**
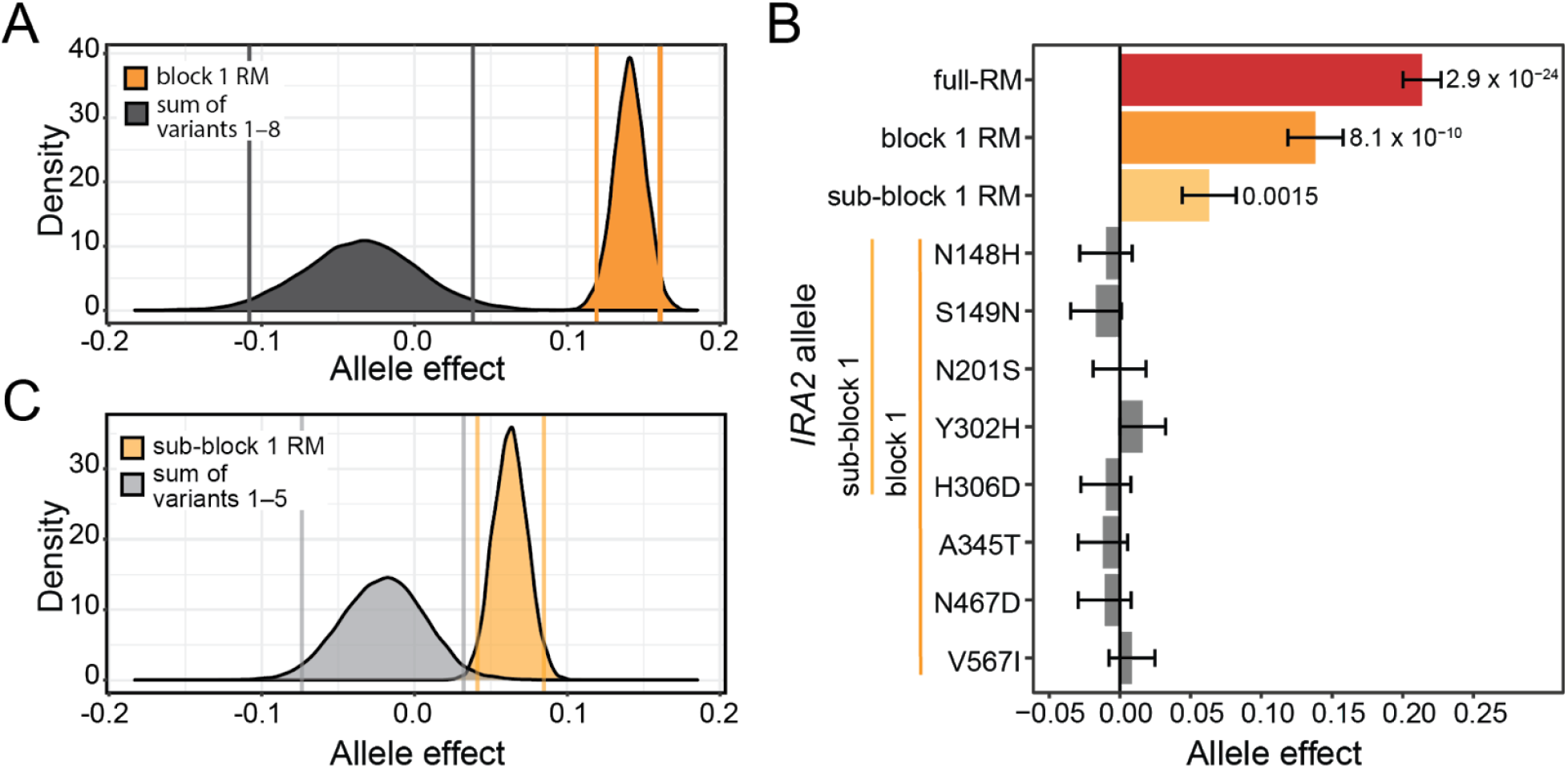
Analysis of the effects of the variants in block 1 and sub-block 1. A: Bootstrap analysis of the effect of the block 1 RM allele compared to the summed effects of the eight variants in block 1. The same-colored vertical lines mark the central 95% quantile range for each distribution. B: Allele effects of the designated *IRA2* genotypes. P-values are for a comparison between the given allele and the BY allele. Only significant p-values less than 0.05 are displayed. Each strain was represented by nine or ten transformants that were each phenotyped three (block 1 RM and sub-block 1 RM) or eight (variants 1–8) times. C: Bootstrap analysis of the effect of the sub-block 1 RM allele compared to the summed effects of the five variants in sub-block 1.

We further divided block 1 into sub-block 1, containing the first five nonsynonymous variants and assayed its effect on Gph1-GFP expression (**Figure 4B; Figure S3 and Table S6**). The effect of sub-block 1 was significant (p = 4 × 10^−7^), but was smaller than the entire effect of block 1 (p = 0.05). This result suggests that the effect of block 1 is caused by multiple variants: at least one among the first five variants and at least one among the following three variants. Furthermore, bootstrap analysis showed that the summed effects of the first five variants are unlikely to account for the effect of sub-block 1 **(Figure 4C)**. Thus, further division of block 1 caused its effects to splinter and uncovered the existence of non-additive interactions among the five variants closest to the 5’-end of the gene.

### No evidence for an effect of synonymous variants in block 1

Instead of epistasis among nonsynonymous variants, a possible alternative explanation for the differences between the large effects we measured for the blocks and the smaller additive effects of their single nonsynonymous variants could be effects caused by the synonymous variants within the blocks. Specifically, the engineered blocks were generated by amplifying the block regions from BY and RM genomic DNA. Therefore, the blocks include synonymous variants that are not present in our single-variant experiments. Although they do not change protein sequences, synonymous variants have been reported to affect complex traits (Sharon et al., 2018; She and Jarosz, 2018). For example, their effects could arise from changes in translation that could affect Ira2 expression or function, especially for variants close to the 5’-end of the gene (Plotkin and Kudla, 2011; Tuller et al., 2010). We chose to test the combined effect of the eleven synonymous variants in block 1. Block 1 had the largest effect of the four blocks, while none of its eight nonsynonymous variants had a significant effect on their own. We engineered alleles of block 1 that carried each of the four different combinations of the synonymous variants and nonsynonymous variants. These alleles showed a strong effect of the nonsynonymous variants in block 1 (ANOVA p = 8 × 10^−7^). The synonymous variants had no discernible effect (p = 0.20; **Figure 5)**. Thus, the overall effect of block 1 must be due to multiple nonsynonymous variants with non-additive interactions.

**Figure 5:**
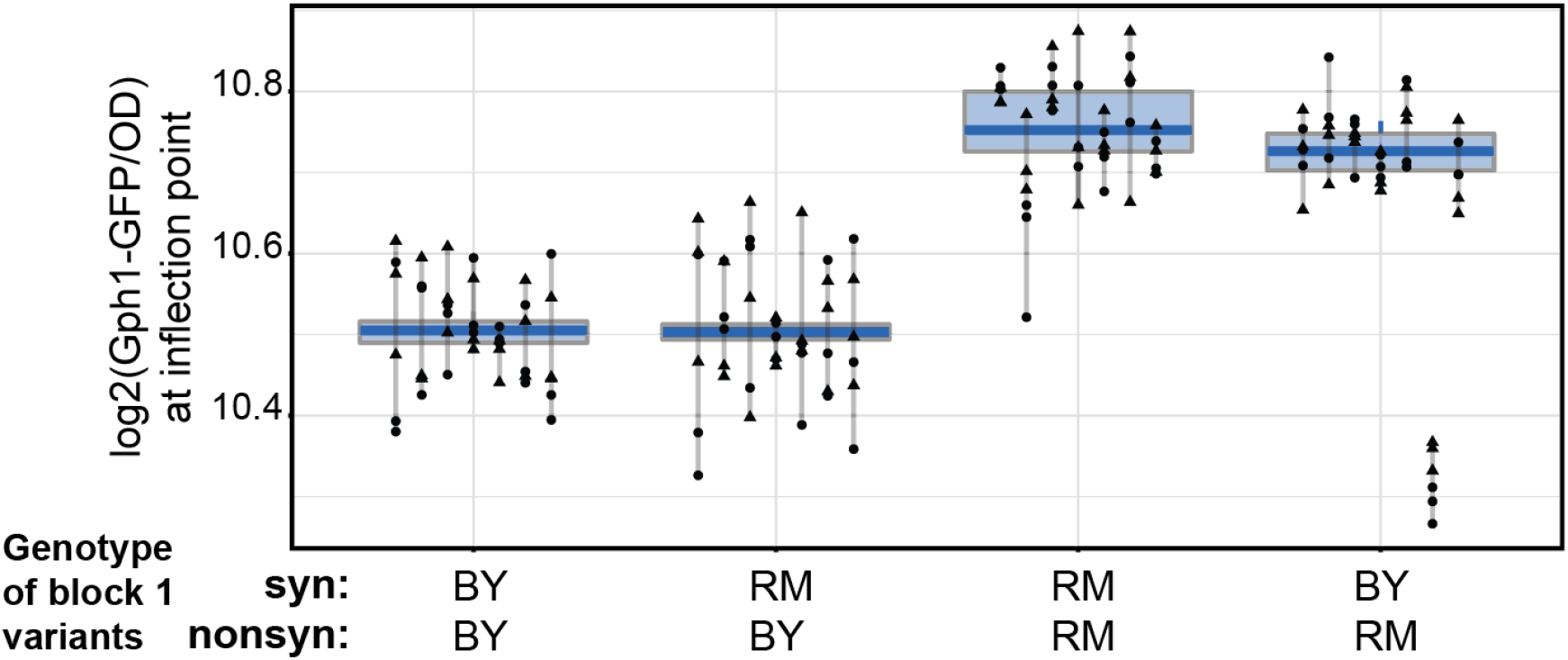
Measurement of the effect of the nonsynonymous variants in block 1. Different shapes represent different plate reader runs. Lines connect measurements of the same transformant. For each strain, seven independent transformants were phenotyped six times. The boxplots show the median as thicker central lines and the first and third quartiles computed on averaged phenotypes per transformant.

## Discussion

Our fine-mapping experiments in the *IRA2* gene revealed multiple causal variants spread throughout the *IRA2* ORF that likely act together to create the *trans*-eQTL hotspot at this gene. Altogether, there must be at least seven causal variants in this gene. In our single-variant experiments, we directly identified one variant each in block 2 and block 3, and two variants in block 4. In addition, there must be at least three causal variants in block 1, with at least two in sub-block 1, that epistatically interact to create the strong increase in Gph1-GFP expression caused by this block. In addition to this localized epistasis within block 1, which covers a poorly conserved region of the ORF with unknown function, we also found evidence for epistasis between variants spread further across the gene body. Thus, this single gene harbors a complex molecular genetic architecture.

We were able to detect the small effects of the causal variants in *IRA2* because in conjunction they create a large effect that manifests as a highly pleiotropic *trans*-eQTL hotspot that drew our attention to this region. It is interesting to speculate how many variants of similar effect may go unnoticed by QTL mapping because they fail to co-occur in a fashion that creates a detectable QTL (Bernstein et al., 2019; Kroymann and Mitchell-Olds, 2005; Metzger and Wittkopp, 2019).

Our work has several limitations. First, we used a single, GFP-tagged gene as a phenotype for fine-mapping *trans* effects of *IRA2*. We cannot rule out that other traits affected by the *IRA2* locus are shaped by different causal variants, possibly in genes close to *IRA2*. However, we deem it unlikely that traits that are truly influenced by the *IRA2* gene are shaped by different variants within *IRA2*, given that Ira2 acts high in the RAS signaling pathway (**Figure 1B**). Assuming that *IRA2* variants ultimately affect phenotypes via altered cAMP/PKA signaling, it is unclear how variants could have trait-specific downstream effects. A possible exception may occur if Ira2 performs unknown functions other than regulating RAS. Second, while we have provided evidence for the existence of epistatic interactions between variants in *IRA2*, we have not determined the specific variants that engage in these interactions. Because epistatic interactions are probably required for explaining the full difference between the BY and RM alleles, the precise molecular basis for this strain difference remains open.

In conclusion, our result of multiple causal variants in the *IRA2* gene mirrors observations about genetic architectures at coarser scales. Quantitative traits including gene expression can be highly polygenic (Bloom et al., 2013; Boyle et al., 2017; Flint and Mackay, 2009; Visscher et al., 2017), and the expression of a given gene is shaped by many loci across the genome (Albert et al., 2018; Brion et al., 2020; Metzger and Wittkopp, 2019). Each of these loci can harbor multiple causal genes (Bernstein et al., 2019; Steinmetz et al., 2002). In turn, these genes can have multiple causal variants, extending the pattern of multifactorial complexity to the finest possible scale of individual nucleotides in a single gene. It remains to be explored whether and why certain genes are more prone to carrying single, strong variants versus multiple variants with smaller effect.

## Materials and Methods

### Yeast strains, primers, and media

The *S. cerevisiae* strains used here are derived from S288C (BY4741 (MATa, his3Δ1 leu2Δ0 met15Δ0 ura3Δ0), referred to as “BY” in the text) and RM-11a, (RM HO(BY) (MATa, his3Δ1::CloNAT, leu2Δ0, ura3Δ0 HO(BY allele) AMN1(BY allele), referred to as “RM”). A complete listing of the strains used in this study can be found in **Supplementary Table 1**. All primers are listed in **Supplementary Table 2**.

We used the following media (recipes are for 1L):

For general yeast strain growth and storage:

YPD (10 g yeast extract, 20 g peptone, 20 g glucose)

For selection of the GFP-HIS3MX6 cassette in *GPH1* tagging:

SDC-His (1.66 g SC -His -Leu -Ura, 100 mg leucine, 200 mg uracil, 20 g glucose)

For selection of the KanMX4 cassette in *IRA2* gene deletions and verifying allele exchanges by CRISPR-Swap:

YPD +G418 (G418 sulfate (Fisher Scientific cat# BP6731); 200 μg/ml)

For selecting for the CRISPR-Swap plasmid in cells after transformation:

SDC-Leu (1.66 g SC -His -Leu -Ura (Sunrise Science; cat# 1327-030), 50 mg histidine, 200 mg uracil, 20 g glucose)

For phenotyping Gph1-GFP expression:

YNB LowFlo (6.7 g yeast nitrogen base -folic acid -riboflavin with ammonium sulfate and without amino acids (Sunrise Science; cat# 1536-050), 20 g glucose, 50 mg histidine, 100 mg leucine, 50 mg methionine, 200 mg uracil). Sterilized by filtration.

For iodine staining:

YNB (6.7 g yeast nitrogen base with ammonium sulfate and without amino acids (BD Biosciences cat# 291940), 20 g glucose, 50 mg histidine, 100 mg leucine, 50 mg methionine, 200 mg uracil)

For solid media, 20 g/L agar was added prior to autoclaving. Yeast were grown at 30°C.

### *GPH1*-GFP tagging and *IRA2* gene deletions

Insertions of cassettes for genome modification were performed using a standard PCR-based one-step method (Longtine et al., 1998). Yeast transformations were performed using a standard LiAc procedure (Gietz and Schiestl, 2007). Transformants expressin g the selectable marker were single colony purified and insertion of the cassette into the correct location and absence of the unmodified wildtype allele were verified by colony PCR.

Strain BY GPH1-GFP:HIS3MX6 (YFA0644) was obtained from the GFP collection (Huh et al., 2003). RM HO(BY) GPH1-GFP:HIS3MX6 (YFA0649) was created as follows: the GPH1-GFP allele was PCR amplified from YFA0644 genomic DNA using primers OFA0471 and OFA0472 and integrated into RM HO(BY) (YFA0254).

Strains BY GPH1-GFP ira2Δ::KanMX (YFA0650 and YFA01430) and RM GPH1-GFP ira2::KanMX (YFA0654) were created by integration of the *ira2*Δ::kanMX allele, which was PCR amplified from YFA0666 genomic DNA using primers OFA0468 and OFA0469. BY GPH1-GFP *ira2*Δ::KanMX was created twice because glycerol stock storage at −80°C of YFA0650 resulted in a stark reduction in CRISPR-Swap efficiency.

Strain BY GPH1-GFP *ira2_block1*Δ::KanMX (YFA1443) was created by replacing *IRA2* block 1 sequence with the KanMX cassette PCR amplified from nej1Δ::KanMX (YFA0007) genomic DNA using primers OFA1079 and OFA1080.

### Creation of *IRA2* allele fragments used as repair templates for CRISPR-Swap

All *IRA2* allele fragments were PCR amplified from genomic DNA or commercially synthesized DNA using Phusion Hot Start Flex DNA polymerase (NEB). Genomic DNA was isolated using the ten-minute preparation (Hoffman and Winston, 1987). All PCR fragments were analyzed by agarose gel electrophoresis, excised, purified using Monarch DNA Gel Extraction Kit (NEB), and quantified using Qubit fluorometric quantification (Thermo Fisher Scientific).

The IRA2(BY) allele was amplified from BY4741 or YLK1879 genomic DNA and the IRA2(RM) allele (sometimes referred to as “full-RM”) from YFA0254 or YLK1950 genomic DNA using primers OFA0468 and OFA0469. These primers amplify a 9,431 bp *IRA2* fragment with termini (123 bp at the 5’ end and 68 bp at the 3’ end) that have identical sequence to the region flanking the *ira2*Δ::KanMX cassette.

The *IRA2*-block alleles were created using gene splicing by overlap extension (SOEing) (Horton et al., 1989). Each of the four block regions was PCR amplified from BY (BY4741 for blocks 1 and 2 and YLK1879 for blocks 3 and 4) and RM (YFA0254) genomic DNA. Primers used were: block 1, OFA0467 and OFA0455; block 2, OFA0456 and OFA0457; block 3, OFA0458 and OFA0459; and block 4, OFA0460 and OFA0544. Pairwise fusions of block 1 and block 2 (4 combinations) were amplified with primers OFA0467 and OFA0457 and block 2 and block 3 (4 combinations) were amplified with primers OFA0458 and OFA0544. Finally, all pairwise fusions of block 1 and 2, and block 3 and 4 (16 combinations) were fused by amplification with primers OFA0468 and OFA0469.

Single-variant alleles were created by PCR SOEing using complementary primers containing the desired RM variant (see **Table S2**). In the first step, two PCR amplifications were performed using BY (YLK1879) genomic DNA as a template. The first amplification used the variant-specific reverse primer and OFA0467, and the second amplification used the variant-specific forward primer and OFA0544. In the second step, the two PCR fragments were fused using primers OFA0468 and OFA0469.

The *IRA2* block1 RM allele was created as described above for the *IRA2*-block alleles. The *IRA2* sub-block 1 RM allele was created by PCR SOEing of two PCR fragments. Fragment 1 was amplified with primers OFA0467 and OFA0589 from RM genomic DNA (YLK1950), and fragment 2 was amplified with primers OFA0536 and OFA0544 from BY genomic DNA (YLK1879). The two PCR fragments were fused using primers OFA0468 and OFA0469.

The *IRA2*-block 1 synonymous and nonsynonymous variant alleles were PCR amplified using primers OFA0468 and OFA1124. Amplification with these primers creates a 1,978 bp *IRA2* fragment with termini (123 bp at the 5’ end and 54 bp at the 3’ end) that are identical to the region flanking the *ira2_block1*Δ::KanMX cassette. The synBYnonsynBY and synRMnonsynRM alleles were amplified from BY and RM genomic DNA (YLK1879 and YLK1950, respectively) and the synBYnonsynRM and synRMnonsynBY alleles were amplified from synthetic DNA (Twist Bioscience).

### CRISPR-Swap

We followed the protocol of (Lutz et al., 2019). For each CRISPR-Swap transformation, the amount of the *IRA2* allele repair fragments ranged from 1000-1500 ng, and the amount of plasmid pFA0055 (Addgene #131774) was 330 ng. pFA0055 expresses Cas9 as well as the guide RNA (gCASS5a) that directs Cas9 to the 5’-end of the KanMX cassette. We recovered 10 to 320 transformants when engineering in BY using the 9,431-kb *IRA2* repair template and 504 to more than 900 transformants when using the 1,978 kb *IRA2* block 1 repair template. Of single colony streaked transformants, 98 – 100% had lost G418 resistance, indicating a successful allele exchange.

The sixteen chimeric block strains were engineered in one batch, as were the strains with the four arrangements of synonymous and nonsynonymous variants in block 1. The BY and RM background strains were engineered in two batches: one created the BY GPH1-GFP IRA2(BY) and RM GPH1-GFP IRA2(RM) strains and the other the BY GPH1-GFP IRA2(RM) and RM GPH1-GFP IRA2(BY) strains. The *IRA2* single-variant strains were engineered in four batches, corresponding to the four *IRA2* blocks. In each batch, a new IRA2(BY) strain was also engineered, and in batches 1, 2 and 4 a new IRA2(RM) strain was engineered. The IRA2(BY) strains from these four batches were similar in their effect on Gph1-GFP expression and therefore were grouped into one IRA2(BY) genotype that served as the wildtype baseline for determining the effects of the other alleles. The IRA2(RM) strains were similarly grouped. The IRA2 block1 RM strain and the IRA2 sub-block 1 RM strain were created in the same batch (block1) as variants 1 – 8.

### Verification of *IRA2* alleles after CRISPR-Swap

All G418 sensitive transformants after CRISPR-Swap were assumed to have exchanged the KanMX allele for the provided *IRA2* allele. To verify the presence of the correct *IRA2* allele and that strain mix-ups did not occur during the experimental procedures, some of the strains were partially genotyped at different stages (see below for details). Verified strains are designated in **Supplementary Table 1**.

The *IRA2* alleles in the BY GPH1-GFP IRA2(BY) (YFA0658 and YFA0659) and RM GPH1-GFP IRA2(RM) (YFA0662 and YFA0663) strains were genotyped as BY or RM based on an EcoRI site present in the RM but not the BY allele. Genomic DNA isolated from these strains was PCR-amplified with OFA0068 and OFA0070. The PCR product was then digested with EcoRI and analyzed by agarose gel electrophoresis.

A representative strain, either transformant #1 or #2, from each of the sixteen *IRA2* block alleles was confirmed to have the expected arrangement of blocks by verification of the expected variants at the block borders after Sanger sequencing. Genomic DNA was isolated from cultures that were started from glycerol stock plates (YPFA010 and YPFA011) and used for PCR amplification. For each strain, the junction between blocks 1 and 2 was amplified using primers OFA0585 and OFA0905 and sequenced with OFA0904, and the junction between blocks 3 and 4 was amplified using primers OFA0906 and OFA0907 and sequenced with OFA0903. Prior to sequencing, the fragments were analyzed by gel electrophoresis and purified using a Monarch DNA Gel Extraction Kit (NEB). All strains had the expected variants in the sequenced fragment. One strain, RRBR #1 had a *de novo* variant in the sequenced region resulting in an aGc to aTc (S1860I) amino acid change. This strain was not removed from phenotyping and its Gph1-GFP expression was similar to that of the other RRBR transformants. None of the RBRR transformants were sequenced.

To identify potential *ira2* loss of function alleles among the transformants of the sixteen different *IRA2* block strains, cells were spotted onto YNB agar plates and stained with iodine vapors.

Iodine stains wildtype yeast cells containing glycogen a dark reddish brown. Cells with elevated RAS-GTP levels, as in the case of an *ira2*Δ, are unable to store glycogen and therefore stain pale yellow in the presence of iodine (Gil and Seeling, 1999). For each of the sixteen strains, 7 – 8 transformants were stained with iodine and 7 / 127 stained pale yellow, possibly due to errors created during the multiple-round PCR amplification of the *IRA2* alleles. Transformants that stained pale yellow were excluded from phenotyping.

In the *IRA2* single-variant strains, the expected amino acid change was verified for transformants of N148H, S149N, N201S, Y302H, H306D, A345T, N467D, V567I, Q757H, and 1507V. Genomic DNA was isolated from cells taken from the randomly arrayed starter culture plates and PCR amplified using primers OFA1053 and OFA1054. The PCR fragment was then Sanger sequenced in the region of the expected variant using the following primers: OFA0592 for N148H and S149N; OFA0589 for N201S; OFA0591 for Y302H, H306D, and A345T; OFA0536 for N467D; OFA0904 for V567I; OFA0539 for Q757H; and OFA0540 for I1507V.

### Phenotyping of Gph1-GFP expression in the plate reader

Starter cultures were inoculated with cells from glycerol stocks and grown overnight at 30°C in 800–1000 μl of YPD medium in a 2-ml deep-96-well-plate. The plates were sealed with a Breathe Easy membrane (Diversified Biotech) and placed on an Eppendorf Mixmate set at 1100 rpm. After overnight growth, the starter culture plates were sealed with aluminum foil and stored at 4°C for the duration of phenotyping.

For each phenotyping plate run, 10 μl of resuspended cells from the starter culture plate was used to inoculate 600 – 800 μl of YNB LowFlo medium in a 2-ml deep-well plate. These pre-cultures were grown overnight as described for the starter cultures. After overnight growth, the pre-cultures were diluted to an OD_600_ = 0.05 in 100 μl of YNB LowFlo medium in a 96-well flat bottom plate (Costar) and the plates were sealed with a Breathe Easy membrane (Diversified Biotech).

The strains were grown for ~24 hours in a Synergy H1 (BioTek Instruments) plate reader at 30°C with readings taken every 15 min for 97 cycles with 10 sec of orbital shaking between reads and 11 – 13 min between cycles. Cell growth was characterized using absorbance readings at 600 nm and Gph1-GFP expression was measured from the bottom of the plate using excitation at 488 nm and emission at 520 nm.

Data was processed as described in (Lutz et al., 2019). Briefly, we fit growth curves to each well to identify the inflection point at which the yeast culture begins to exit exponential growth. We extracted the fluorescence and OD_600_ values at this point as well as the two time points flanking it, took their respective averages, and calculated the log_2_ of the ratio between the average fluorescence value and the average OD_600_ value as the Gph1-GFP expression level for the given well.

### Designation of *IRA2* domains

Conserved regions between Ira2 and NF1 were determined by multiple sequence alignment of Ira2/Ira1/NF1 sequences from *Candida glabrata* (KTB00138.1), *Saccharomyces paradoxus* (translated from CP020290.1) *Kluyveromyces lactis* (translated from CR382125.1), *Homo sapiens* (P21359.2) and *Saccharomyces cerevisiae* (AAA34710.1 and AAA34709.1) using Clustal Omega (Sievers et al., 2011), as provided through the EMBL-EBI analysis tool API (Madeira et al., 2019).

The CHD (Neurofibromin CTD-homology domain) and Sec-PH (Sec14 homologous and pleckstrin homology-like domain) were defined by previous studies (D’angelo et al., 2006; Luo et al., 2014). We defined here the Threonine-Serine Rich Domain (TSRD). The TSRD extends from amino acid 399 to 1021 and is enriched for serines and threonines, but not enriched for cysteines. This region has 23 / 24 of the detected phosphorylation sites in Ira2 (Holt et al., 2009; Lanz et al., 2021; Swaney et al., 2013). We chose the start of the TSRD at a cluster of serines and the end at the last detected phosphorylated residue. The TSRD is 16.5% serine and 10.6% threonine, while the flanking amino acids 1 – 398 and 1022 – 1645 have 13.1% and 6.3% serine and 8.5% and 5.3% threonine residues, respectively.

### Ancestral alleles and BY and RM allele frequencies

Ancestral alleles were determined by comparison to two different evolutionary outgroups to BY and RM. First, we determined the nucleotide allele present in the closely related species *S. paradoxus* after an alignment of *IRA2*(BY) to GenBank #AABY01000044.1. Second, we obtained the nucleotide allele present in the Taiwanese *S. cerevisiae* isolate EN14S01 (‘standardized name’: ‘AMH’) from (Peter et al., 2018). EN14S01 is a member of the highly diverged clade 17 thought to have branched early from all other *S. cerevisiae* isolates. Ancestral alleles defined by these two outgroups are in good agreement (Table 1). The population allele frequencies of the BY and RM variants across 1,011 *S. cerevisiae* species were obtained from (Peter et al., 2018). Predictions of variant effects were obtained using PROVEAN (Choi and Chan, 2015) (http://provean.jcvi.org) and are based on amino acid conservation and properties. Variant changes resulting in scores of less than −2.5 are predicted to be deleterious.

### Statistical analyses of allele effects

All statistical analyses were conducted in R (https://www.r-project.org) version 4.0.4. The effect of RM alleles on the expression of GFP-Gph1 (*y*) was estimated using the “lmer” function in the lme4 package (Bates et al., 2015) for fitting mixed-effects linear models. In each case, one RM allele (such as a block or a single variant, see below) was compared to strains with the BY allele that were measured during the same runs in the plate reader. The models included the genotype (BY vs. the given RM allele) as a fixed effect, plate and transformant as random effects (denoted in parentheses), and the residual error (*ε*):

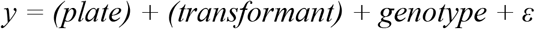

The effects of genotype were estimated as slopes of the linear models. Significance was calculated by using type I ANOVAs, as provided by the lmerTest package (Kuznetsova et al., 2017). These linear models and ANOVAs were performed to test the following effects:

1. The RM vs BY allele in both the BY or RM background
2. The various BY / RM combinations of the four blocks
3. The effect of each individual nonsynonymous variant compared to BY

### Testing for epistasis among the four “block” regions of *IRA2*

We tested for epistasis between blocks with a linear model. The model fitted the effects of all four blocks simultaneously, along with all two-way, three-way, and the four-way interactions:

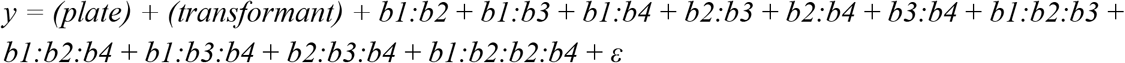

We tested the significance of each term using a Type I ANOVA through sequential addition of terms to the model above.

### Testing for epistasis between single nucleotide variant effects

The single nucleotide variants were not engineered in all combinations, precluding the use of interaction terms in a linear model. Instead, we asked whether the effects of single variants could sum to the observed effect of their given multi-variant block or the full *IRA2* RM allele. To properly take into account measurement error across our experimental design, in which individual transformants were run multiple times in several different runs of a plate reader, we used bootstrapping. For each set of variants, we performed stratified sampling with replacement across measurement plates, ensuring that each genotype was represented at the same sample size as in the full data. For each bootstrapped experiment, we fit the same models as above and extracted the effect estimate. Single-variant effect estimates were summed to calculate the effect they would be expected to have together under an additive model. We performed 10,000 of these bootstraps to generate two distributions. The first distribution represented the difference between BY and the given multi-variant RM allele, and the second distribution represented the sum of the respective single-variant effects. Significance testing was performed by calculating the overlap between the two distributions. If the central 95% quantile range of the two distributions did not overlap, we considered it significantly unlikely that the additive effects of the single variants are able to account for the observed effect of the multi-variant allele. Thus, the less the two bootstrapped distributions overlap, the stronger the evidence for epistatic interactions among the given set of single variants.

## Supporting information

All Supplementary Tables

## Code availability

All analysis code is available at: https://github.com/Krivand/Multiple-causal-DNA-variants-in-a-single-gene-affect-gene-expression-in-trans

## Strain availability

All strains constructed in this work are available on request.

## Supplementary Materials

**Figures**

Figure S1: Gph1-GFP expression in chimeric *IRA2* block alleles

Figure S2: Gph1-GFP expression in single-variant alleles

Figure S3: Gph1-GFP expression in block 1 alleles

**Tables**

Table S1: Strains

Table S2: Primers/Oligonucleotides

Table S3: Allele effects and linear models of full RM in BY or RM background

Table S4: Allele effects and linear models of all block combinations compared to BY

Table S5: Allele effects and linear models of single RM variants

Supplementary Table 6: Allele effects and linear models of block 1 RM, sub-block 1 RM, and their single variants

## Acknowledgements

We thank Laura Sherer, Kaushik Renganaath, Laura Johnson, and Christian Brion for technical contributions and discussions.

## Funding

This work was supported by NIH grant R35GM124676 to FWA.

---

**Figure S1.**
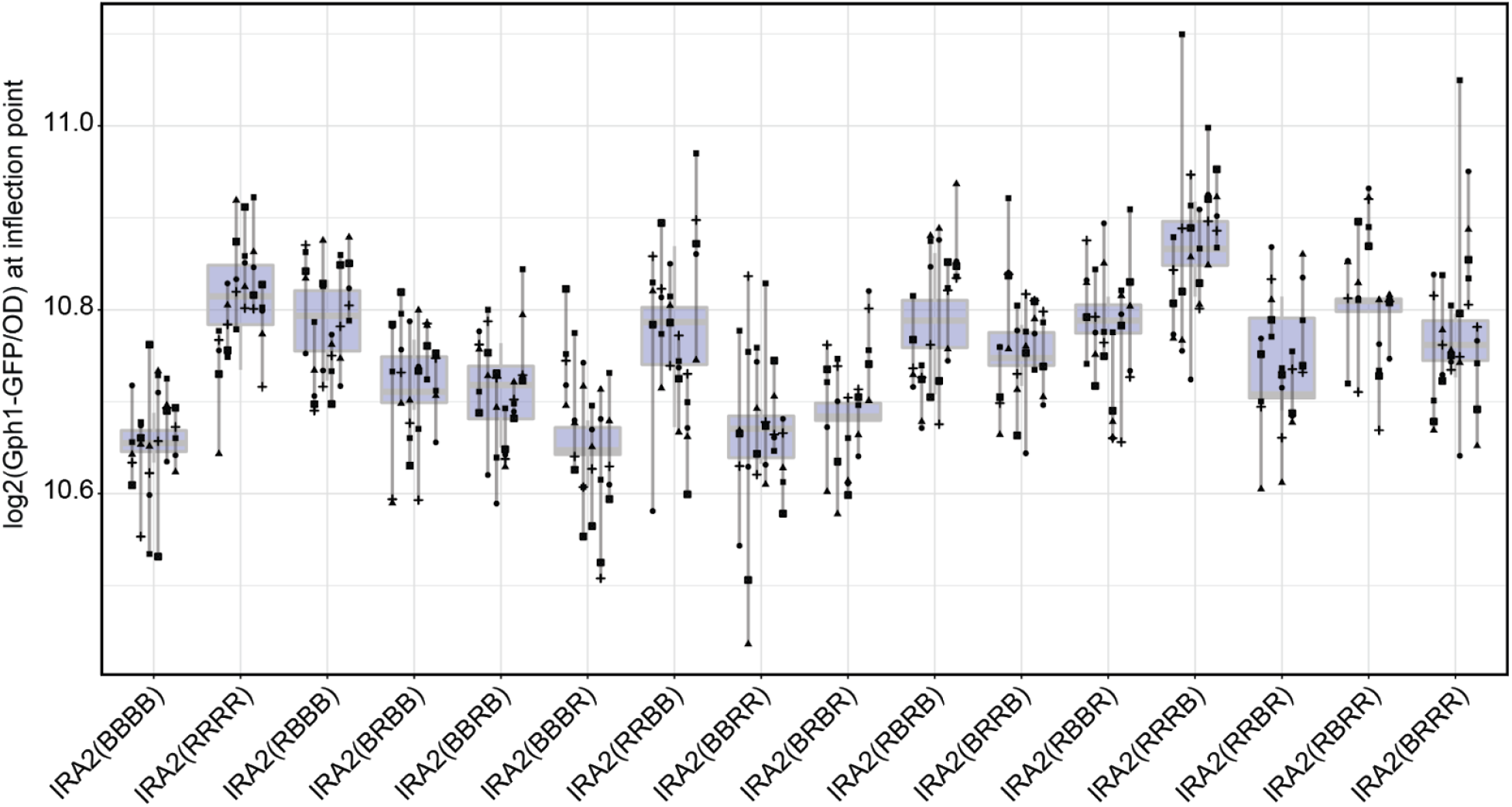
Measurement of Gph1-GFP expression in each of the 16 chimeric *IRA2* block alleles. Chimera alleles are indicated as four letters that are either B or R, indicating the BY or RM allele, respectively, at blocks 1 through 4. Different shapes represent different plate reader runs. Lines connect measurements of the same transformant. For each strain, five or six independent transformants were phenotyped six times. The boxplots show the median as thicker central lines and the first and third quartiles computed on averaged phenotypes per transformant.

**Figure S2.**
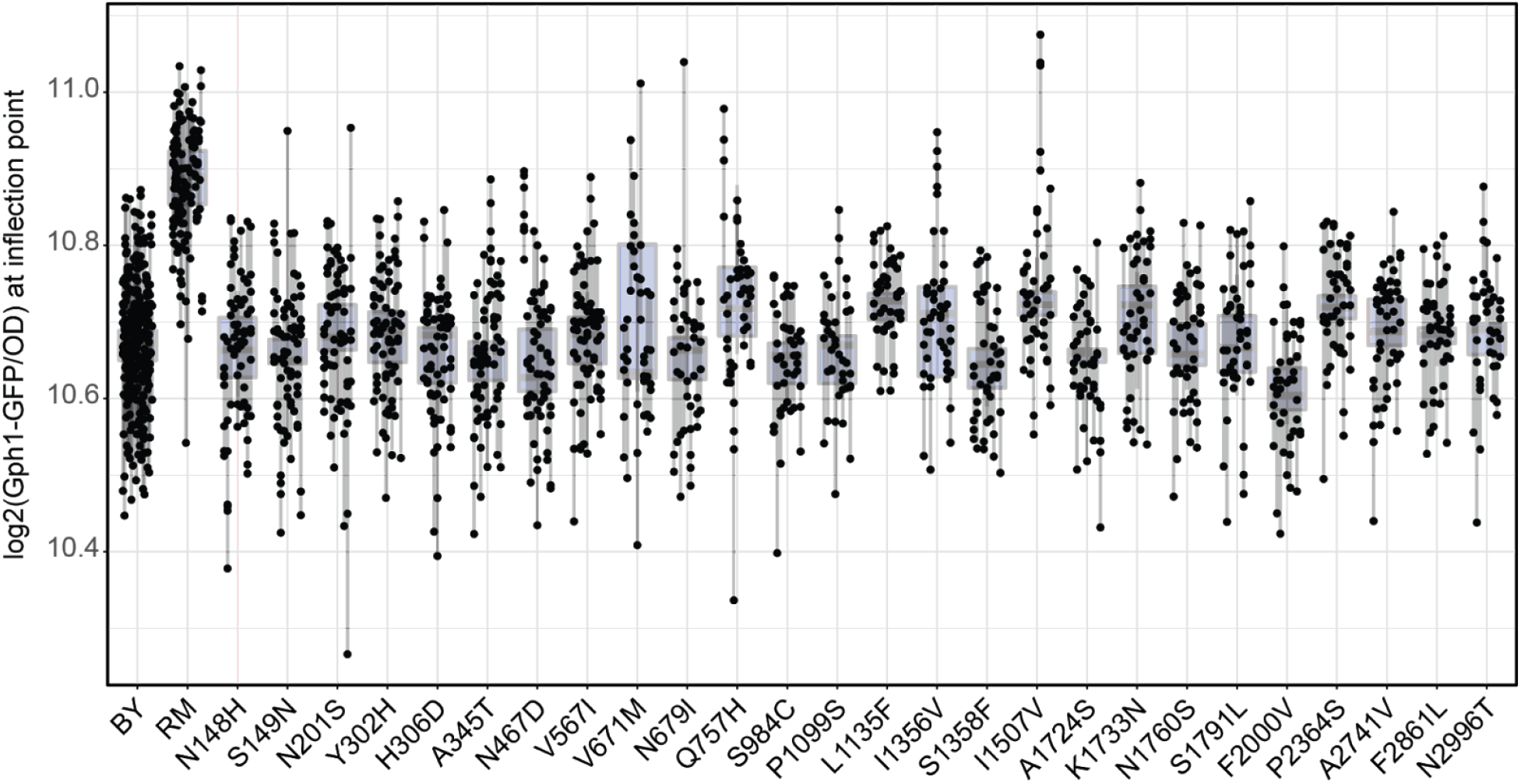
Measurement of Gph1-GFP expression in each single-variant strain. Lines connect measurements of the same transformant. The *IRA2*(BY) allele is represented by 36 transformants, each measured 8 times. The *IRA2*(RM) allele is represented by 27 transformants, 9 of which were measured 8 times and 18 of which were measured 3 times. Single-variant alleles 1–8 are represented by 9 different transformants, each measured 8 times. Single-variant alleles 9–26 are represented by 9 transformants, each measured 5 times. The boxplots show the median as thicker central lines and the first and third quartiles computed on averaged phenotypes per transformant.

**Figure S3.**
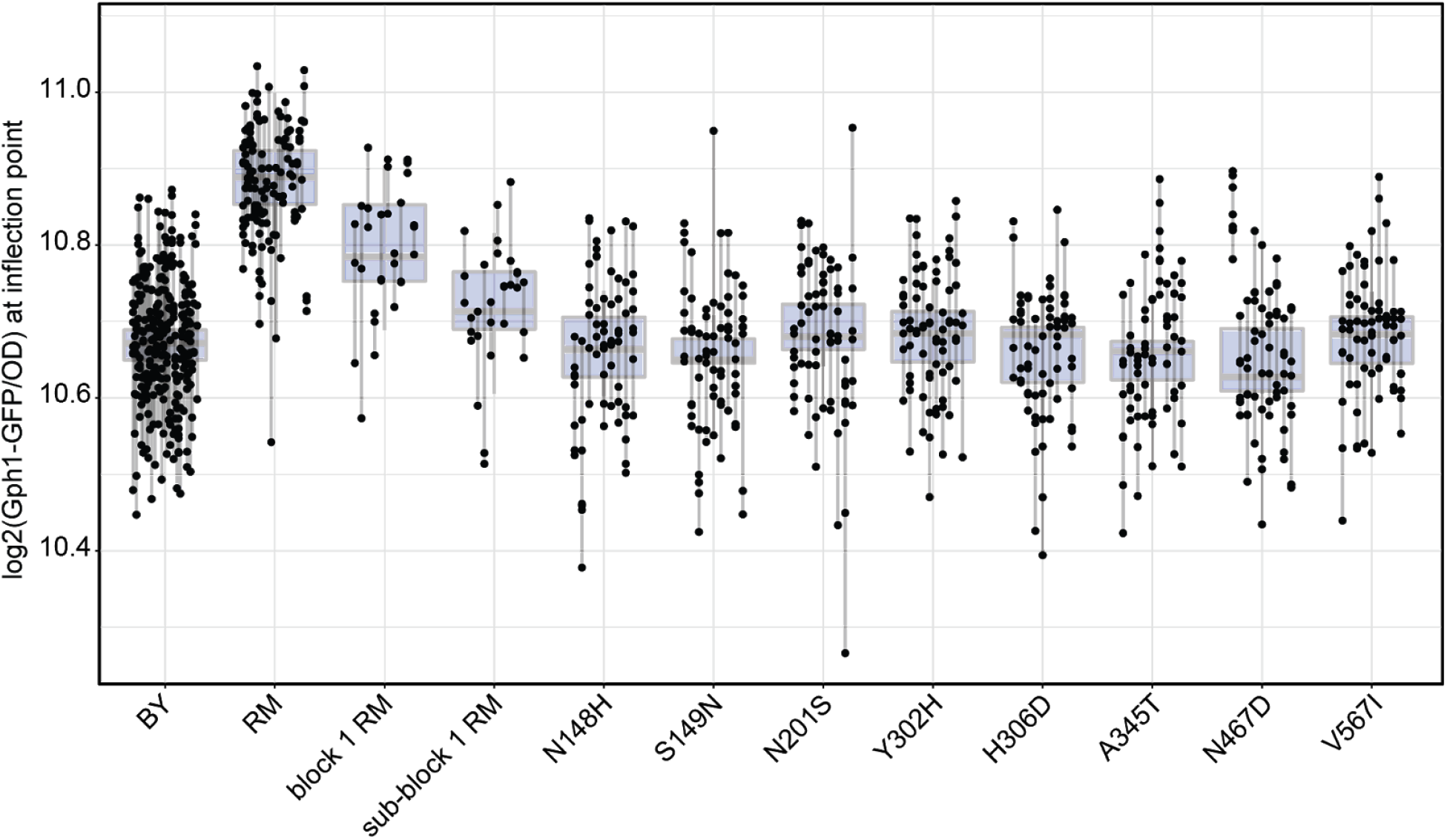
Measurement of Gph1-GFP expression of block1 variants, alone and in combination. Lines connect measurements of the same transformant. The *IRA2*(BY) allele is represented by 36 transformants, each measured 8 times. The *IRA2*(RM) allele is represented by 27 transformants, 9 of which were measured 8 times and 18 of which were measured 3 times. Single-variant alleles 1–8 are represented by 9 different transformants, each measured 8 times. The block 1 and sub-block 1 alleles are each represented by 10 transformants measured 3 times. The boxplots show the median as thicker central lines and the first and third quartiles computed on averaged phenotypes per transformant.

## Notes

### Competing Interest Statement

The authors have declared no competing interest.

